## Supplemental text, figures and tables for "A Hepatitis C virus genotype 1b post-transplant isolate with high replication efficiency in cell culture and its adaptation to infectious virus production in vitro and in vivo"

**This PDF file includes:**

Supplementary Text
Figs. S1 to S10
Table S1

**Other Supplementary Materials for this manuscript include the following:**

Movies S1 to S5

### Supplementary Text

#### Supplementary Methods

##### PHH

Primary human hepatocytes were prepared in house at Heidelberg University Hospital. The human liver tissue used to extract PHHs was obtained from a patient undergoing liver cirrhosis which was approved by the French National Ethics Committee and legal instances. No patient information was available in the laboratory. After extraction, PHH were plated in 6-well culture dishes at $1.5x10^6$ cells/well. Protein lysates were prepared 4 days after seeding.

##### Western blot

For SDS-PAGE/Western blotting, approximately $1\times10^6$ cells were lysed in 50 µl lysis buffer (50 mM Tris-HCl, pH 7.4, 150 mM NaCl, 15 mM $MgCl_2$, 1% Triton X-100) containing an EDTA-free protease inhibitor cocktail pill (Roche) for 1 h on ice. After centrifugation with 14,000 rpm for 30 min at 4 °C, the cleared supernatant was mixed with 50 µl 2×Laemmli buffer and denatured at 95 °C for 10 min. The lysate of approximately $1\times10^5$ cells was loaded in one lane of a polyacrylamide–SDS gel with an appropriate percentage for the protein of interest. The color-prestained protein standard, broad range (11 to 245 kDa) (New England Biolabs), was used to determine the apparent molecular weight of the proteins. After the separation, proteins were transferred to a PVDF membrane using a semi-dry blotter according to the instructions of the manufacturer. The membrane was blocked for 1 h in 5% milk or BSA in 0.5% TBS-Tween20. The primary antibody was diluted in 3% milk-0.5% TBS-T and incubated overnight at 4 °C shaking. After three washing steps with 0.5% TBS-T, the secondary antibody was diluted in 3% BSA or milk in 0.5% TBS-T for 1 h at room temperature. The detection was done using Clarity ECL blotting substrate (Bio-Rad) and the Advanced ECL imaging system (Intas Science Imaging Instruments). The signal intensity was quantified using Fiji.

##### Immunofluorescence

Specific staining of PI4P on intracellular membranes has been described elsewhere (*80*). In brief, for expression of the HCV NS3-5B proteins, $4\times10^4$ Huh7-Lunet T7 cells were seeded 24 h prior to transfection in a 24-well plate on coverslips. Transfection was done with LT1 transfection reagent (Mirus Bio LLC, Madison, WI, USA) according to the manufacturer's instructions. Cells were fixed 18 h post-transfection in 4% PFA for 20 min at room temperature followed by permeabilization with 0.5% Digitonin-PBS for 20 min. The permeabilized cells were then blocked for 1 h in 5% BSA-PBS prior to incubation with the primary antibodies in 3% BSA-PBS for 1 h at room temperature. After three washing steps with PBS each for 5 min, the secondary antibodies (Alexa Flour, Invitrogen) were incubated in 3% BSA–PBS for 45 min in the dark at room temperature. After three washing steps with PBS each for 5 min, nuclei were stained with a 1:4,000 TBS dilution of 4′,6-diamidino-2-phenylindole (DAPI) for 1 min. Cells were mounted with Fluoromount G (Southern Biotechnology Associates), and images were acquired with a Leica SP8 confocal laser-scanning microscope (Leica Microsystems). Quantification of cellular PI4P levels was performed as previously described (*17*). Briefly, whole-cell z-stacks of the PI4P channel were recorded as 8-bit TIFF files using 20 slices of around 0.4 µm thickness with a ×40 (numerical aperture (NA) 1.4) objective of at least three different randomly chosen fields of view. For each stack, a single slice was recorded for the NS5A channel to identify NS5A-positive cells for PI4P quantification. For quantification of NS5A signals, stacks were recorded in an identical manner as for PI4P. Image stacks were z-projected according to their maximum intensity in Fiji and a constant threshold was set to 30–40, creating a binary image. The NS5A channel was overlaid, NS5A-positive cells were encircled and integrated density values of the thresholded PI4P signal were acquired by Fiji software.

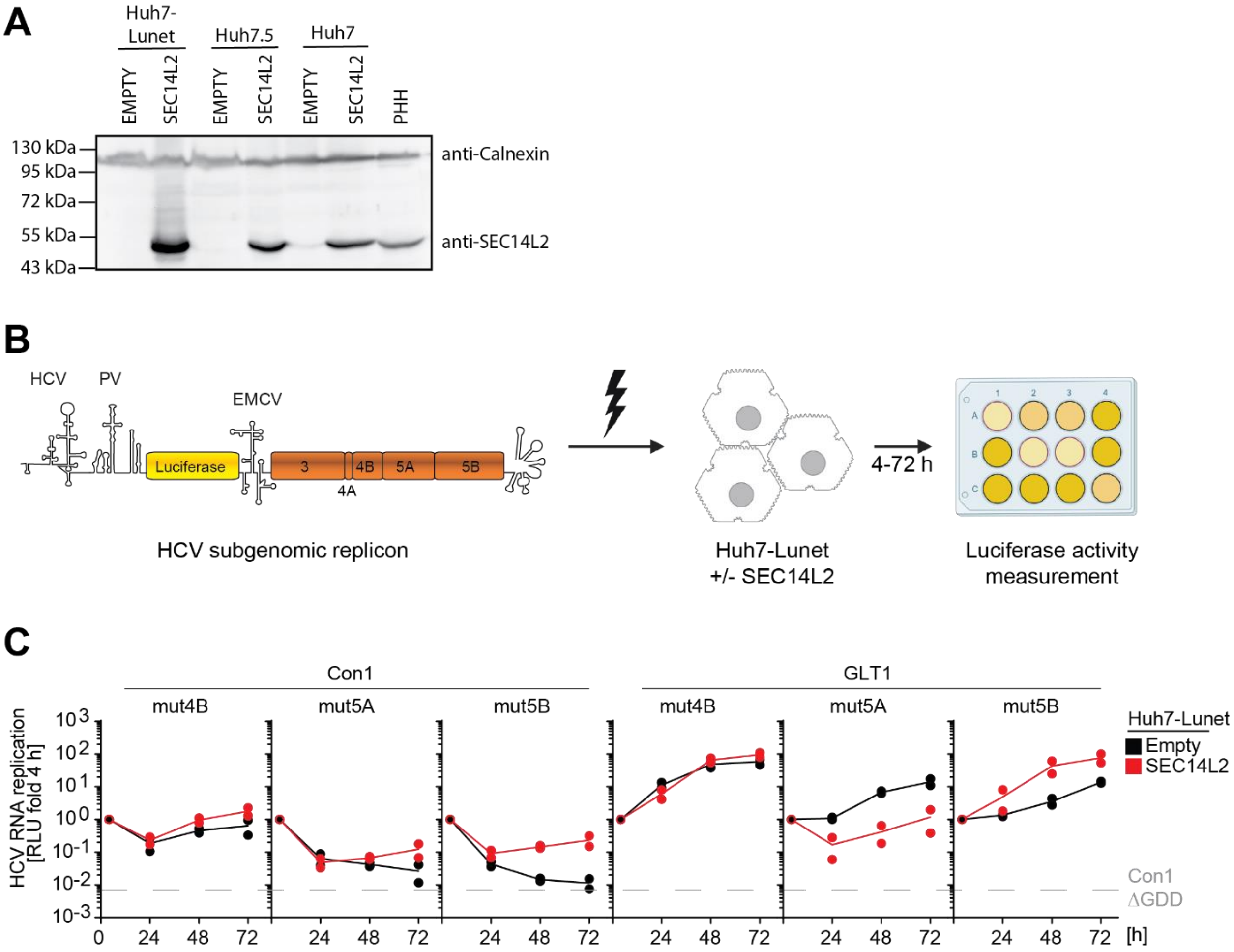


**Fig. S1: SEC14L2 expression, schematic of the experimental procedure and replication efficiency of GLT1 in Huh7-Lunet cells.** (**A**) SEC14L2 expression levels in Huh7-Lunet, Huh7.5 and Huh7 cells after lentiviral transduction with a SEC14L2 encoding vector or empty control, compared to PHH. Approximately $1x10^5$ Huh7-Lunet, Huh7.5, Huh7 or primary human hepatocytes (PHH) were lysed and analyzed by 10% SDS-PAGE/Western blotting for SEC14L2 and Calnexin expression. One representative experiment from two independent repetitions is shown. (**B**) Schematic of a subgenomic reporter replicon and of the experimental procedure of luciferase-based replication measurement. HCV 5'UTR (HCV) poliovirus internal ribosomal entry site (IRES) (PV) and EMCV-IRES (EMCV), as well as the HCV 3'UTR are indicated by their respective secondary structures. NS3-NS5B and firefly luciferase coding regions are indicated by orange and yellow boxes, respectively. Transfection by electroporation is visualized by a lightning-symbol. (**C**) Replication enhancement by a combination of mutations and SEC14L2 expression for Con1 (left) and GLT1 (right). Huh7-Lunet cells either expressing SEC14L2 or transduced with empty vector were electroporated with the indicated subgenomic reporter replicon RNA containing mutations in NS4B (K1846T), NS5A (S2204R) or NS5B (R2884G) as indicated. Luciferase activity in cell lysates (RLU) was quantified as a correlate of RNA replication efficiency at the given time points and normalized to 4 h. A replication deficient Con1 variant (Con1ΔGDD) was used as a negative control and the respective luciferase level at 72 h is indicated by a dashed grey line in all diagrams. The data are the mean values from two independent experiments shown as individual data points with two technical replicates each.

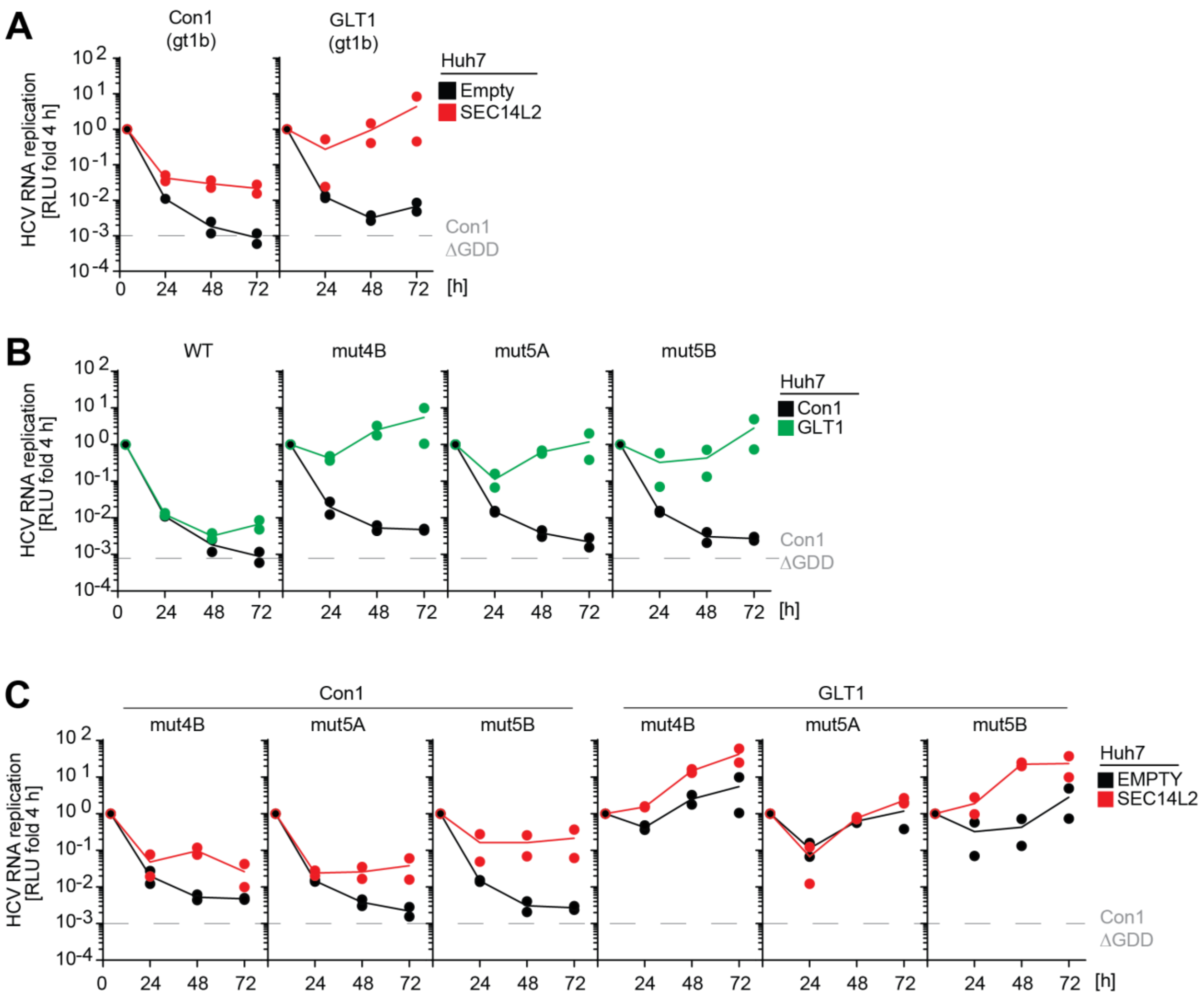


**Fig. S2: Replication efficiency of GLT1 compared to Con1 in Huh7 cells, using different replication enhancing conditions.** Huh7 cells were transfected with subgenomic reporter replicons of the indicated isolates or mutants. Luciferase activity in cell lysates (RLU) was quantified as a correlate of RNA replication efficiency at the given time points and normalized to 4 h. (**A,C**) HCV replication was stimulated by SEC14L2 expression compared to empty vector transduction as indicated. (**B,C**) Replication enhancement of GLT1 (green lines) or Con1 (black lines) by mutations in NS4B (K1846T), NS5A (S2204R) or NS5B (R2884G). A replication deficient Con1 variant (Con1ΔGDD) was used as a negative control for replication and the respective luciferase level at 72 h is indicated by a dashed grey line in all diagrams. The data are the mean values from two independent experiments shown as individual data points with two technical replicates each.

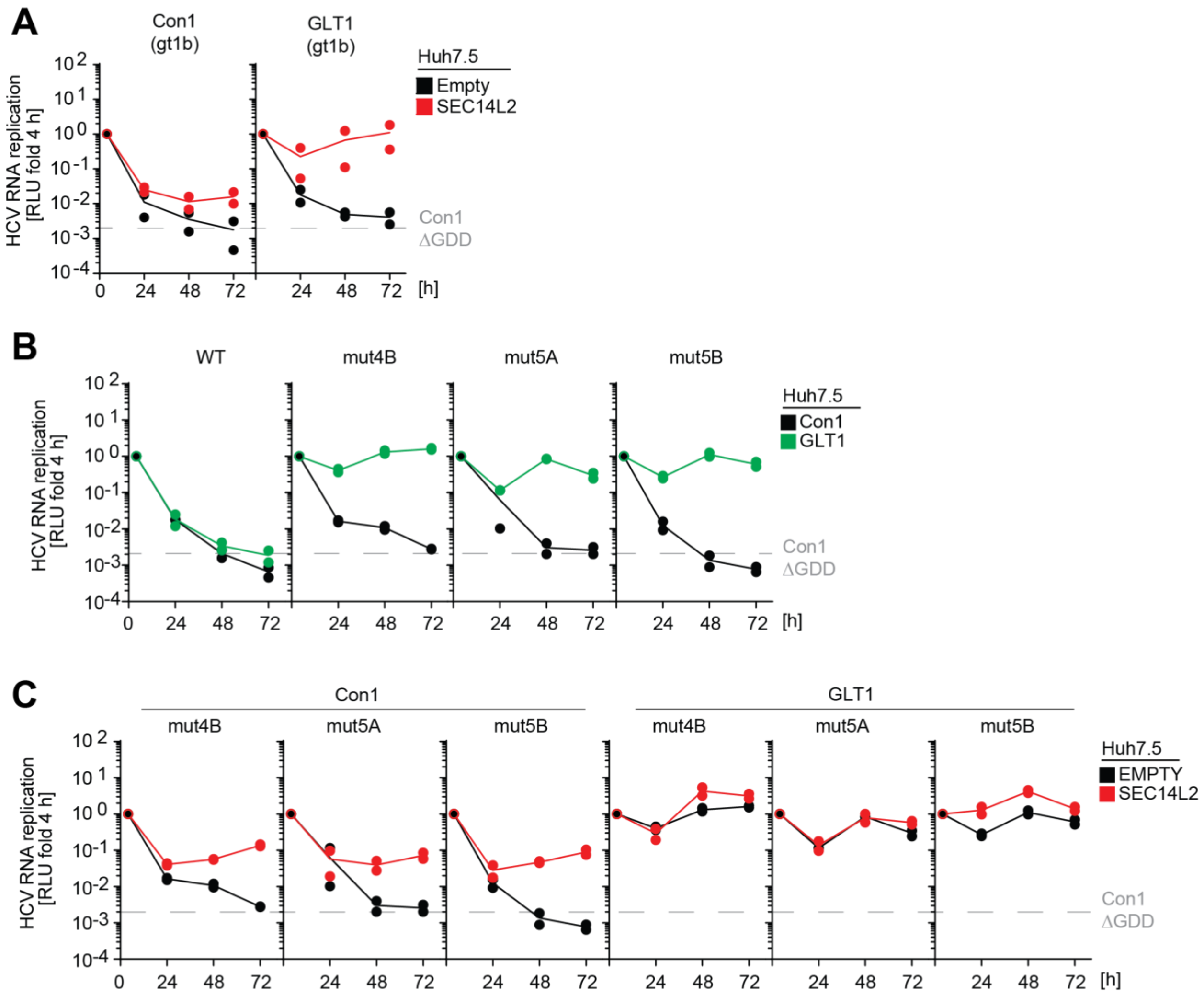


**Fig. S3: Replication efficiency of GLT1 compared to Con1 in Huh7.5 cells, using different replication enhancing conditions.** Huh7.5 cells were transfected with subgenomic reporter replicons of the indicated isolates or mutants. Luciferase activity in cell lysates (RLU) was quantified as a correlate of RNA replication efficiency at the given time points and normalized to 4 h. (**A,C**) HCV replication was stimulated by SEC14L2 expression compared to empty vector transduction as indicated. (**B,C**) Replication enhancement of GLT1 (green lines) or Con1 (black lines) by mutations in NS4B (K1846T), NS5A (S2204R) or NS5B (R2884G). A replication deficient Con1 variant (Con1ΔGDD) was used as a negative control for replication and the respective luciferase level at 72 h is indicated by a dashed grey line in all diagrams. The data are the mean values from two independent experiments shown as individual data points with two technical replicates each.

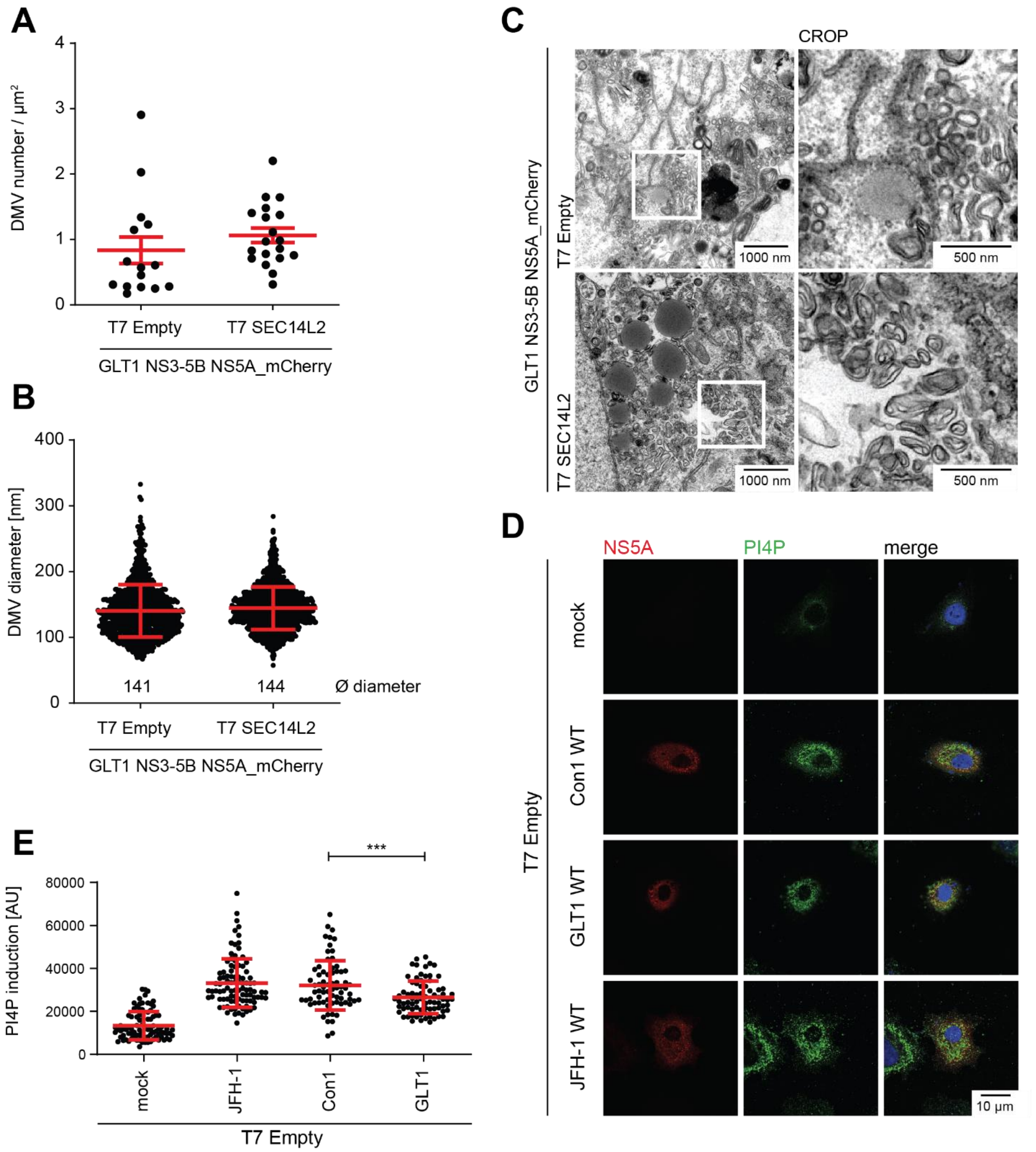


**Fig. S4: Ultrastructural analysis of membrane rearrangements after expression of GLT1 NS3-5B NS5A_mCherry and PI4KA activation by different isolates.** Huh7-Lunet T7 cells either expressing SEC14L2 (T7 SEC14L2) or not (T7 empty) were transfected with a pTM vector encoding GLT1 NS3-5B with an in frame insertion of mCherry in domain 3 of NS5A (*68*). Cells were fixed 24 hours post transfection and the mCherry fluorescent signal was used to identify positive cells for further analysis by CLEM. A minimum of six cells were analyzed and the (**A**) number of DMVs in randomly selected areas and (**B**) the DMV diameter was measured. (**C**) Representative images of each condition. (**D,E**) Huh7-Lunet T7 cells were transfected with plasmids encoding NS3-5B of Con1 WT, GLT1 WT or JFH-1 WT and fixed 24 h post transfection. (**D**) Total NS5A and PI4P was detected by immunofluorescence analysis using monoclonal antibody 9E10 (red) and anti-PI4P (green), respectively. The nuclei were stained with DAPI. (**E**) Quantification of the PI4P signal intensity in 71-95 cells using Fiji. Each dot represents one cell. Results shown are mean values from two independent experiments. Unpaired two-tailed Student's t-test was used to determine statistical significance (*** = $p \leq 0.001$).

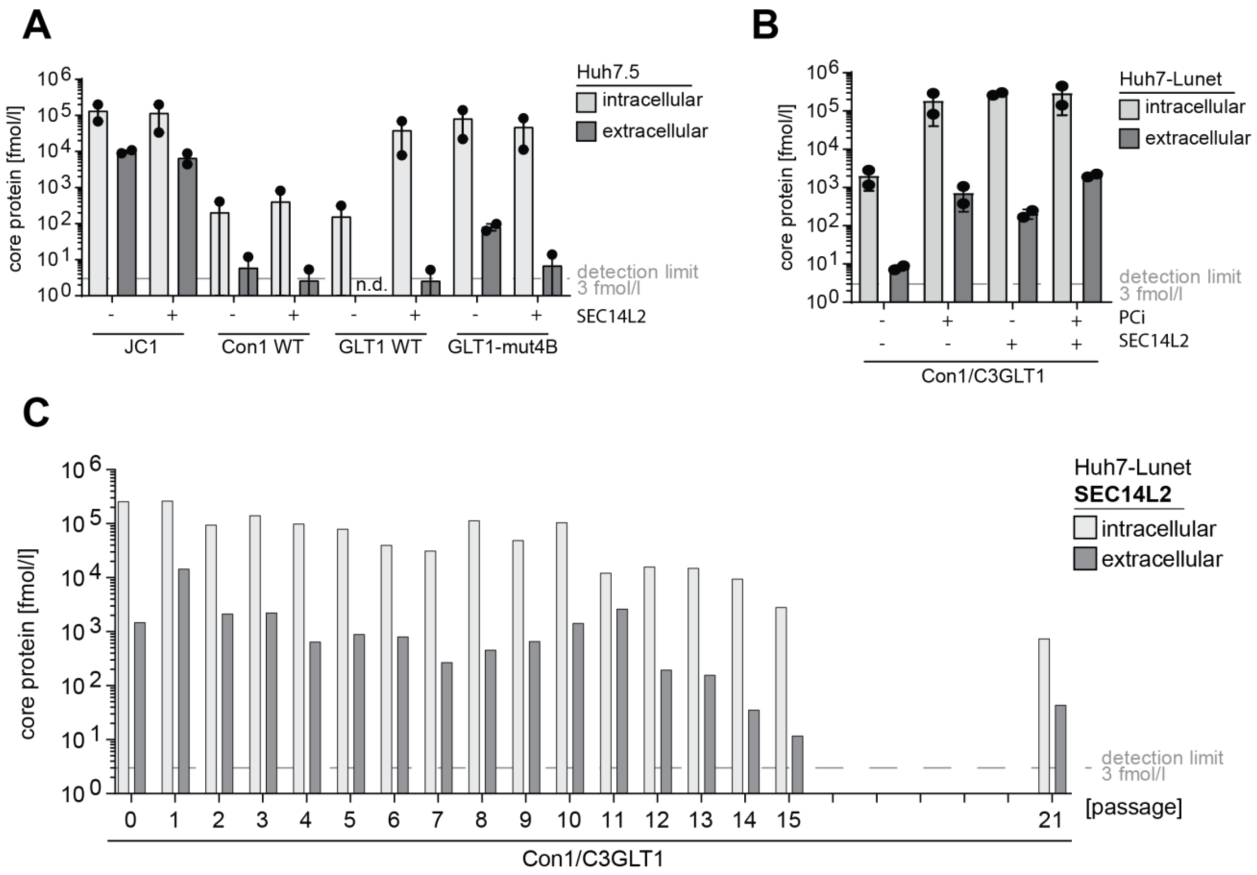


**Fig. S5: Virus production in Huh7.5 cells and analysis of a chimeric GLT1 genome harboring the structural proteins of Con1.** (**A**) Detection of intra- and extracellular core protein after transfection of full-length virus genomes. Huh7.5 cells with or without SEC14L2 expression were transfected with the indicated HCV full-length genomes and intra- and extracellular core protein levels were determined by ELISA as correlates of replication and virus secretion, respectively. (**B,C**) A chimeric GLT1 genome, encoding the structural proteins of Con1 up to the C3 junction site in NS2 (*24*) was transfected in Huh7-Lunet (B) or Huh7-Lunet CD81 (C) cells with or without SEC14L2 expression and/or PCi treatment. Intra- and extracellular core levels were determined by Elisa 72 hours post transfection (B) or before passaging (C). Shown are data from two independent experiments (A, B) or from a single passaging experiment (C). The dashed grey line indicates the detection limit of 3 fmol/l core protein; n.d. = not detectable.

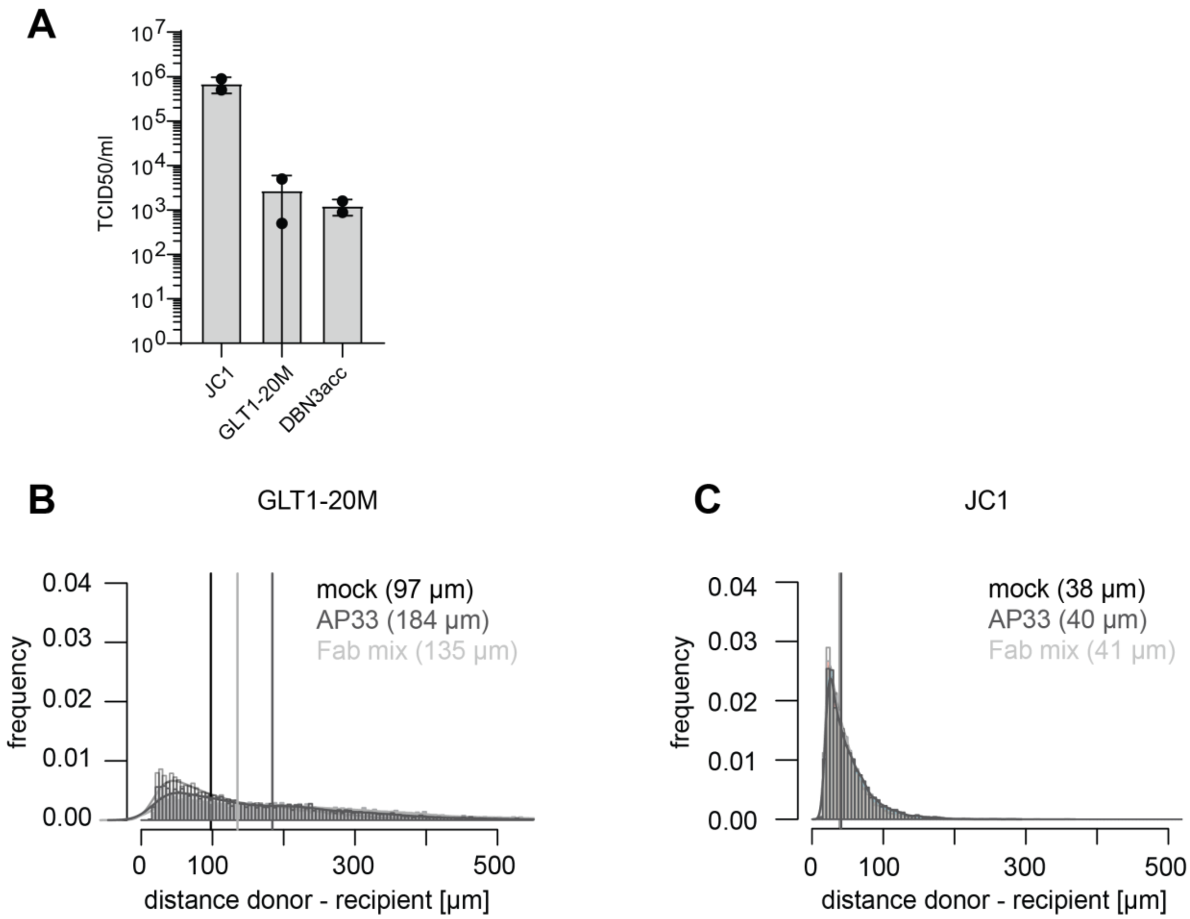


**Fig. S6: Input viral titers and analysis of spreading properties of GLT1-20M compared to JC1.** (**A**) Viral titer (TCID50/ml) from concentrated supernatant of different HCV isolates harvested from Huh7-Lunet CD81 MAVS-GFP-NLS used in Fig. 4D. (**B,C**) Distribution of the shortest distances between a GLT1-20M (B) or JC1 (C) positive Huh7-Lunet CD81 MAVS-GFP-NLS cell ("donor") and the next nearest Huh7-Lunet CD81 MAVS-mCherry-NLS with nuclear mCherry signal ("recipient") after treatment with HCV neutralizing antibodies or left untreated (mock). Numbers in brackets and vertical lines indicate the median distance for each treatment. Shown are representative data from one experiment (n=2).

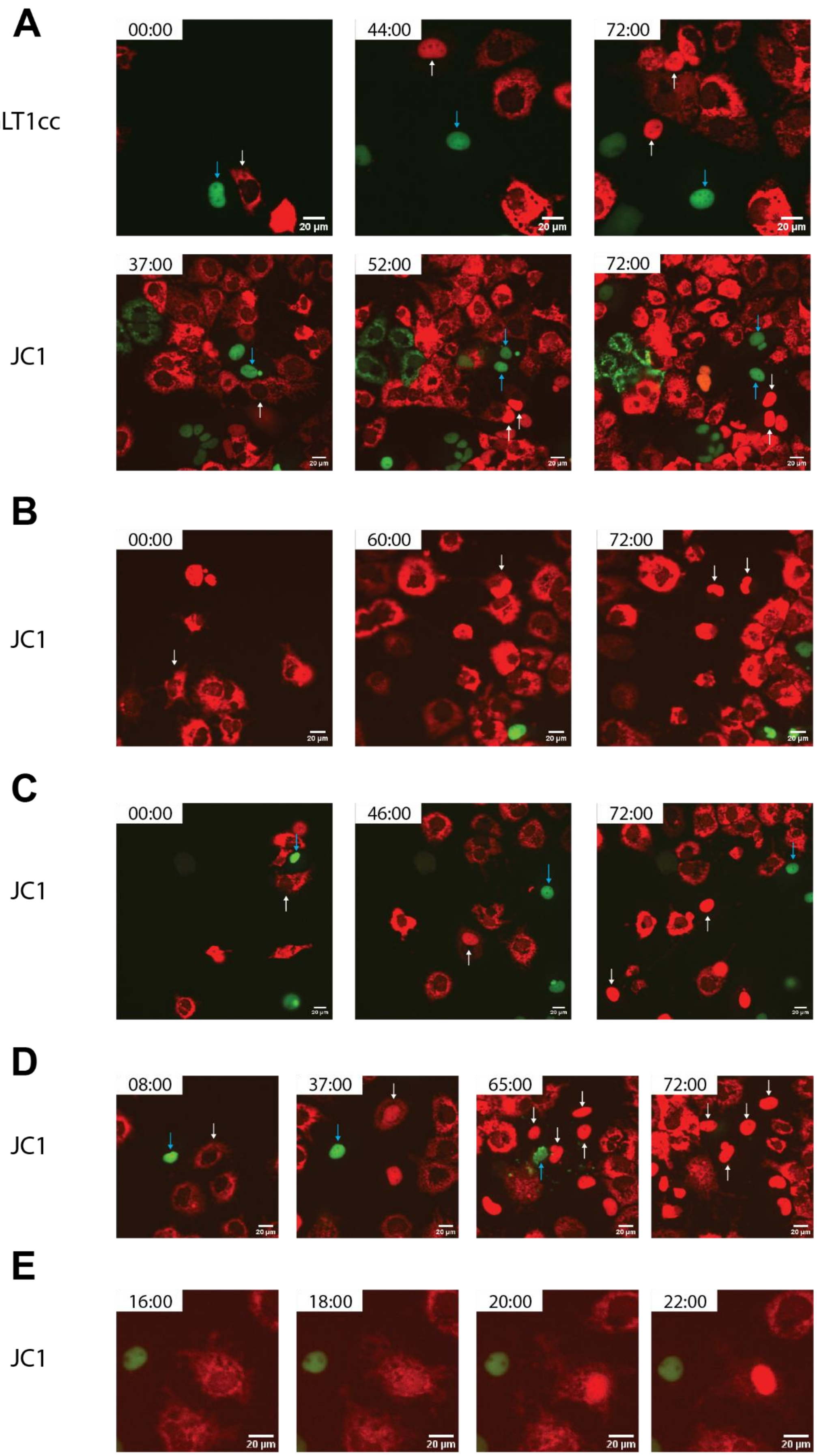
A
GLT1cc
00:00
44:00
72:00
JC1
37:00
52:00
72:00
B
JC1
00:00
60:00
72:00
C
JC1
00:00
46:00
72:00
D
JC1
08:00
37:00
65:00
72:00
E
JC1
16:00
18:00
20:00
22:00
20 µm

**Fig. S7: Representative still images from live cell imaging of virus spread assays.** Every 30 minutes an image was acquired for 72 hours. Donor cells of interest are labelled with a cyan arrow, recipient cells of interest are labelled with a white arrow. (**A**) Example of cell-to-cell mediated spread in both GLT1cc (Movie S1) and JC1 (Movie S2). (**B-E**) Examples based on JC1 because of the overall higher number of infection events. (**B**) Infection in absence of a visible donor cell representing cell-free spread (Movie S3). (**C**) Cell-to-cell spread with recipient cell moving away from donor cell (Movie S4). (**D**) Vanishing of donor cell after cell-to-cell spread (Movie S5). (**E**) Earliest observed infection event.

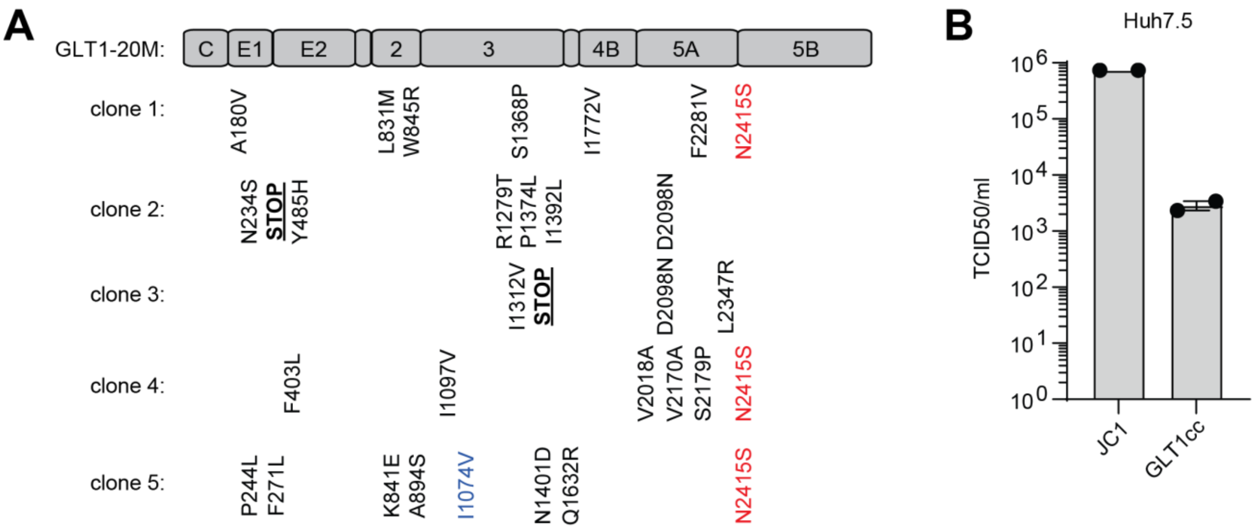


**Fig. S8: Identification of polymorphic sequence N2415S in the quasispecies of p118.27 and titer of GLT1cc in Huh7.5 cells.** (**A**) Sequences of 5 individual subclones of RT-PCR products obtained from total RNA of p118.27 compared to GLT1-20M. A missing consensus mutation at position 1074 is shown in blue, premature stop codons are underlined and highlighted in bold. N2415S is shown in red color. (**B**) Viral titer (TCID50/ml) from concentrated supernatant of indicated HCV isolates harvested from Huh7.5 cells; data from two independent biological replicates.

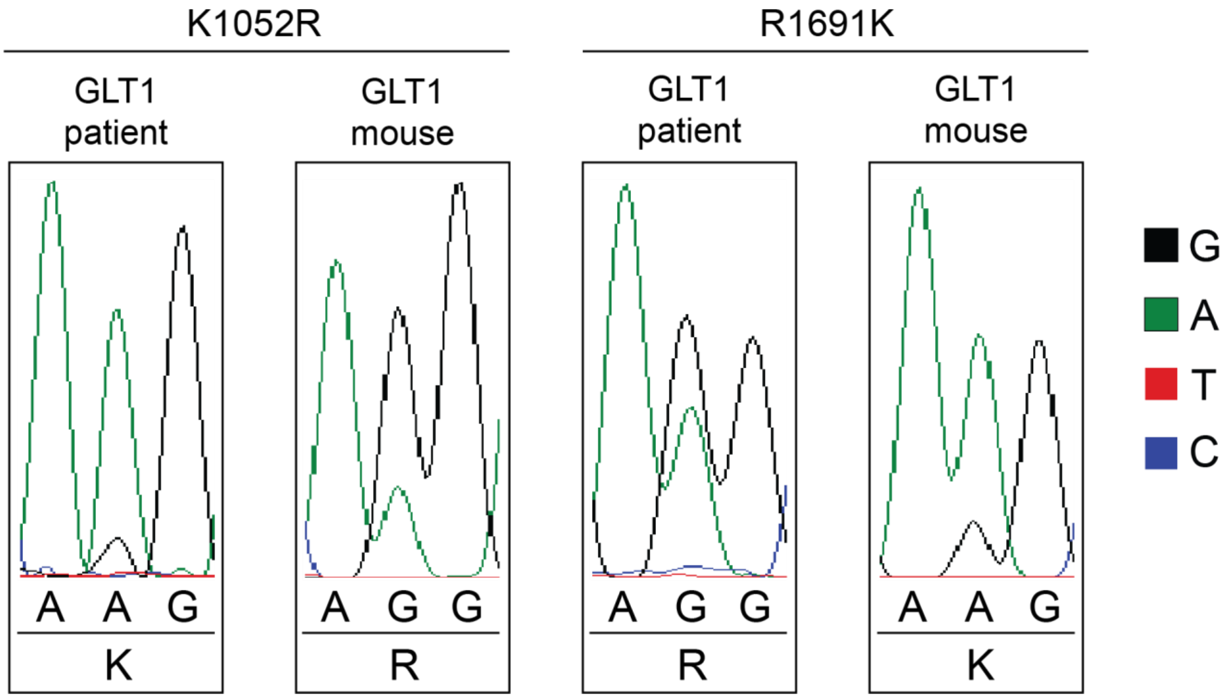


**Fig. S9: Sequence chromatograms of individual base positions with a shift in abundance in the GLT1-patient serum compared to the mouse serum 8 weeks after infection.**

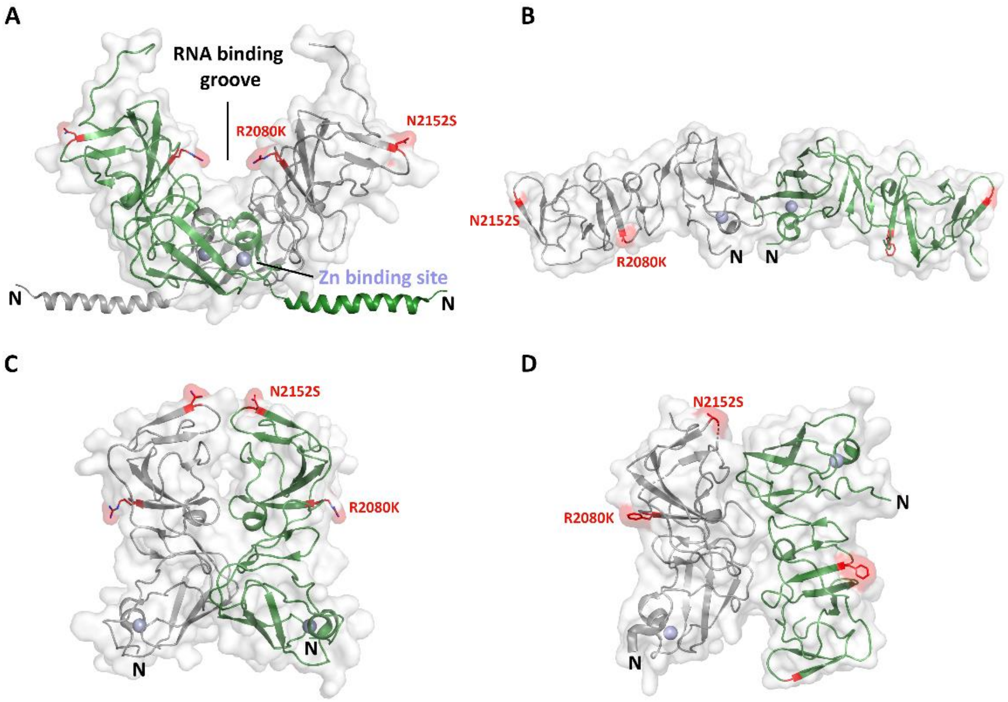


**Fig. S10: Alternative NS5A domain 1 dimer structures.** The NS5A domain 1 in is anchored to phospholipid membranes by an N-terminal amphipathic helix (residues 1–33, PDB 1R7G). Crystal studies revealed four different dimeric forms of domain 1 from genotype 1a and 1b with the same monomeric unit, but different dimeric arrangements shown. (**A**) The first crystal structure from genotype 1b (PDB: 1ZH1) with the modelled N-terminal amphipathic helix and the postulated RNA-binding groove between the two monomers is shown. (**B**) Monomers from genotype 1a (PDB: 4CL1) form a head to head dimer. (**C**) Genotype 1b monomers assemble in parallel to form an extensive interface (PDB: 3FQM). (**D**) Monomers from genotype 1a dimerize via a similar interface as shown in C but are assembled in an antiparallel fashion (PDB: 4CL1).

**Table S1: DNA oligonucleotides used in this study.**

| Oligo Name | Sequence | Use |
|---|---|---|
| A_9416 | CAGGATGGCCTATTGGCCTGGAG | cDNA generation |
| S_59 | TGTCTTCACGCAGAAAGCGTCTAG | cDNA amplification/qPCR |
| A_3580 | GTCCAGCACACGCCATTGAC | cDNA amplification |
| S_93 | AAGCGTCTAGCCATGGCGTT | cDNA amplification |
| A_3489 | GGCCTGTGAGGCTAGTGATGATACA | cDNA amplification |
| S_3317 | GTGCGGGGACATCATCTCGG | cDNA amplification |
| A_6103 | GCTATCAGCCGGTTCATCCACTGC | cDNA amplification |
| S_3420 | TCGCCCATCACGGCCTAC | cDNA amplification |
| A_5982 | TGACCAGGTCCTCGGTGGAG | cDNA amplification |
| S_4540 | GTAACGAGCTCGCCGCGCAGCTGTC | cDNA amplification |
| A_9386 | TTAGCTCCCCGTTCATCGGTTGG | cDNA amplification |
| S_5120 | GTGCGCCAGGGCTCAGGCTCCACC | cDNA amplification |
| A_9364 | GGAGCAGGTAGATGCCTACCCCTAC | cDNA amplification |
| GLT1_probe | 6-FAM–TCCTGGAGGCTGCACGACACTCAT–TAMRA | qPCR |
| A_165 | TACTCACCGGTTCCGCAGA | qPCR |
| Jc1_probe | 6-FAM–AAAGGACCCAGTCTTCCCGGCAATT–TAMRA | qPCR |
| S_146 | TCTGCGGAACCGGTGAGTA | qPCR |
| A_219 | GGGCATAGAGTGGGTTTATCCA | qPCR |
| pFK_bb_fwd | TTCAACGACTCCATGGCCTTAGCGCATTTTC | pFK i341 PiLuc NS3-3´ GLT1-20M, pFK i341 PiLuc NS3-3´ GLT1cc |
| pFK_bb_rev | TGTGAGGCTAGTGATGATACAGCTAAGCATGC | pFK i341 PiLuc NS3-3´ GLT1-20M, pFK i341 PiLuc NS3-3´ GLT1cc |
| GLT1cc_fwd | GTATCATCACTAGCCTCACAGGCCGGGA | pFK i341 PiLuc NS3-3´ GLT1-20M, pFK i341 PiLuc NS3-3´ GLT1cc |
| GLT1cc_rev | AAGGCCATGGAGTCGTTGAATGATCTGAGGTAGGTC | pFK i341 PiLuc NS3-3´ GLT1-20M, pFK i341 PiLuc NS3-3´ GLT1cc |
| S_JCN2a | CTGTGGTGGTTGTGCTATCTCC | pFK i389 GLT1/C3JFH-1 N2A, pFK i389 GLT1cc/C3JFH-1 N2A |
| A_JCN2a | GGGCCCGGGATTTTCCTC | pFK i389 GLT1/C3JFH-1 N2A, pFK i389 GLT1cc/C3JFH-1 N2A |
| S_GLT1_Insert | GAAAATCCCGGGCCCATGAGCACGAATCCTAAACCTC | pFK i389 GLT1/C3JFH-1 N2A, pFK i389 GLT1cc/C3JFH-1 N2A |
| A_GLT1_Insert | GCACAACCACCACAGGAGCTTCGCGAGGAACACTTTATAG | pFK i389 GLT1/C3JFH-1 N2A, pFK i389 GLT1cc/C3JFH-1 N2A |
| S_EcoRI | ACGCATTCTGGCGGAATTCAG | pFK i389 JcN2AΔE1E2 |
| A_NotI | CAGAATATAGTGACGGCCCACG | pFK i389 JcN2AΔE1E2 |
| A_E1 | CTTCTCTAGTGCAGCGGAGACCG | pFK i389 JcN2AΔE1E2 |
| S_E2 | CGGTCTCCGCTGCACTAGAGAAG | pFK i389 JcN2AΔE1E2 |

| | | |
|---|---|---|
| S_KpnI | CAAGCTTGGTACCGAGCTCGGATCCA TGGACCTCATGGGGTA | pcDNAΔcE1E2-GLT1, pcDNAΔcE1E2-GLT1cc |
| A_XbaI | TAGGGCCCTCTAGATTAGGCCTCAGC CTGGGCT | pcDNAΔcE1E2-GLT1, pcDNAΔcE1E2-GLT1cc |
| S_437_mut | GAACTGGGTTCCTTGCCGC | pFK GLT1cc L437F |
| A_437_mut | GCGGCAAGGAACCCAGTTC | pFK GLT1cc L437F |
| S_ClaI | GTAAGGTTATCGATACCCTCACAT | pFK GLT1cc L437F |
| A_XhoI | CGCTCTCCTCGAGTCCAATTG | pFK GLT1cc L437F |
| S_GLT1_K1846T | TAGGTCTTGGGACGGTGCTTGTGGAC | pFK i341 PiLuc NS3-3´ GLT1 K1846T (mut4B), pFK GLT1-mut4B |
| A_GLT1_K1846T | GTCCACAAGCACCGTCCCAAGACCTA | pFK i341 PiLuc NS3-3´ GLT1 K1846T (mut4B), pFK GLT1-mut4B |
| S_GLT1_S2201R | GCCAGCTCTTCCGCTCGCCAGTTGTC | pFK i341 PiLuc NS3-3´ GLT1 S2201R (mut5A) |
| A_GLT1_S2201R | GACAACTGGCGAGCGGAAGAGCTGG C | pFK i341 PiLuc NS3-3´ GLT1 S2201R (mut5A) |
| S_GLT1_R2884G | CATTCAAGGACTCCATGGCCTTAG | pFK i341 PiLuc NS3-3´ GLT1 R2884G (mut5B) |
| A_GLT1_R2884G | CTAAGGCCATGGAGTCCTTGAATG | pFK i341 PiLuc NS3-3´ GLT1 R2884G (mut5B) |
| S_GLT1_NS5A_mcherry | CAAGCGGTCCGAAGGGGAGCCG | pTM NS3-5B GLT1-NS5A_mCherry |
| A_GLT1_NS5A_mcherry | CAAGCGGTCCGAAGGGGAGCCGG | pTM NS3-5B GLT1-NS5A_mCherry |
| S_N2415S | GGTGATAGTGTAGTCTGCTGCTC | pFK-GLT1-mut4B+N2415S, pFK-GLT1-20M+N2415S (GLT1-21M/GLT1cc) |
| A_N2415S | GAGCAGCAGACTACACTATCACC | pFK-GLT1-mut4B+N2415S, pFK-GLT1-20M+N2415S (GLT1-21M/GLT1cc) |
| S_1905 | TTCGGCGTCCCTACGTACAGTTG | pFK Con1/C3GLT1 |
| A_Con1_MMC3 | CTCAGCTCTGGTGATAAGATACTGTA ACCACCATATGAGCCTAGCGAG | pFK Con1/C3GLT1 |
| S_MMC3 | CAGTATCTTATCACCAGAGCTGAGG | pFK Con1/C3GLT1 |
| A_3254 | AACTGCCACCGCAAGGTCTCGTAG | pFK Con1/C3GLT1 |

**Movies S1-S5: Live cell imaging of cell spread assay.** Every 30 minutes an image was acquired for 72 hours. Donor cells were labelled with GFP and recipient cells with mCherry.

**Movie S1: Example of cell-to-cell mediated spread in GLT1cc.**

**Movie S2: Example of cell-to-cell mediated spread in JC1.**

**Movie S3: Infection in absence of a visible donor cell representing cell-free spread in JC1.**

**Movie S4: Cell-to-cell spread with acceptor cell moving away from donor cell in JC1.**

**Movie S5: Vanishing of donor cell after cell-to-cell spread in JC1.**
